# DLPFC Transcriptome Defines Two Molecular Subtypes of Schizophrenia

**DOI:** 10.1101/116699

**Authors:** C. Harker Rhodes, Elijah F. W. Bowen, Jack L. Burgess, Richard Granger

## Abstract

Little is known about the molecular pathogenesis of schizophrenia, possibly because of unrecognized heterogeneity in diagnosed patient populations. We analyzed gene expression data collected from the dorsolateral prefrontal cortex (DLPFC) of post-mortem frozen brains of 189 adult diagnosed schizophrenics and 206 matched controls. Transcripts from 633 genes are differentially expressed in the DLPFC of schizophrenics as compared to controls at Bonferroni-corrected significance levels. Seventeen of those genes are differentially expressed at very high significance levels (< 10^−8^ after Bonferroni correction).

Weighted Gene Co-expression Network Analysis (WGCNA) of the schizophrenic subjects, based on the transcripts differentially expressed in the schizophrenics as compared to controls, divides them into two groups: "Type 1" schizophrenics, have a DLPFC transcriptome similar to that of controls with no expressed genes identified in this subcohort while the "type 2" schizophrenics have a DLPFC transcriptome dramatically different from that of controls, with 3,652 expression array probes to 3,200 genes detecting transcripts that are differentially expressed at very high significance levels. These findings were re-tested and replicated in a separate independent cohort, using the RNAseq data from the DLPFC of an independent set of schizophrenics and control subjects.

We suggest the hypothesis that these striking differences in DLPFC transcriptomes, identified and replicated in two populations, imply a fundamental biologic difference between these two groups of patients who have been diagnosed as schizophrenic.

## Introduction

In spite of decades of work using every available anatomic, histologic, and molecular technique, little progress has been made elucidating the pathobiology of schizophrenia. In many ways the situation has not changed very much since Fred Plum called schizophrenia “the graveyard of neuropathologists” almost a half century ago [1]. A popular hypothesis to explain this lack of progress is that schizophrenia is a heterogeneous disease and that meaningful results have been obscured in studies which pool data from biologically different patients.

There are two publicly available sources of molecular data which can be used to test that hypothesis. The first dataset was generated by scientists in the Clinical Brain Disorders Branch of the Intramural Research Program at NIMH, under the direction of Dr. Daniel Weinberger; it consists of Illumina HumanHT-12 v4 expression array data from the dorsolateral prefrontal cortex (DLPFC) of postmortem brains of almost a thousand patients with psychiatric disease (including schizophrenia and other diagnoses) and neurologically normal matched controls. (Although those investigators have never published their analysis of that data, the data itself is publically available through dbGaP; Study ID: phs000979.) The second relevant dataset contains RNAseq data from postmortem DLPFC collected by the Common Mind Consortium and made publicly available through their website [2].

We report on an analysis of the NIMH expression array data and observe that the schizophrenics in that cohort are of two types. The “type 1” patients have a DLPFC transcriptome very similar to that of the controls while the “type 2” patients have a dramatically different DLPFC transcriptome with several thousand genes differentially expressed compared to the controls. We then replicated that observation in the CMC dataset.

## Methods

### Sources of data

Over a period of many years, and at great effort and expense, the Clinical Brain Disorders Branch of the NIMH intramural program assembled a large collection of frozen human brains from Medical Examiner patients and conducted detailed post-mortem psychiatric reviews to establish their diagnoses. The human tissue collection and processing protocols have been previously described [3, 4]. Poly-A RNA was prepared from dorsolateral prefrontal cortex and hippocampus. Illumina HumanHT-12 v4 expression array data was generated according to the manufacturer’s protocols, and that data was made publicly available at dbGaP (Study ID: phs000979).

With the appropriate IRB approval and dbGaP authorization, that data was downloaded to a secure high-performance cluster (two Intel Xeon x5660 2.8GHz CPUs with 6 cores each, with hyperthreading yielding 24 virtual cores, with 72GB of DDR3-1333 RAM). Data analysis done using Rstudio and protocols for reproducible research. HTML-formatted R markdown files which contain the computer code for the entire analysis are included as supplemental material.

The data used in the replication phase of this study is from the CommonMind Consortium (CMC; http://www.synapse.org/ CMC), a large collaborative group which collected RNAseq data from the DLPFC of schizophrenics and controls. The details of the tissue collection and data generation is described in the primary paper reporting that work [2].

### Pre-processing of the NIMH expression array data

The decrypted dbGaP data includes a directory “PhenotypeFiles” containing multiple files with the subject annotation data from this study. That information was re-formatted as a data frame whose rows are the study subjects and whose columns contain information about the subject (supplemental R code #1).

The decrypted dbGaP data also includes a directory “ExpressionFiles” which contains the expression array data formatted as idat files. Using the Bioconductor package {beadarray}, that data was quantile normalized, log2-transformed, and formatted as a matrix whose rows are the study subjects and whose columns are the expression array data for each probe. At the same time the function calculateDetection(){beadarray} which implements Illumina’s method for calculating the detection scores was used to create a second matrix with those scores (supplemental R code #2).

The expression array dataset initially contained data from 48,107 Illumina probes. It was filtered to remove data from:

1. 2414 probes for which the log2-transformed data was “NA” or “Inf” for any of the subjects;
2. 33,158 or 73% of the probes where, based on the Illumina detection score, the level of expression was statistically significant in fewer than 841 of the 849 subjects;
3. 652 probes where the probe sequence contains a common SNP [5].

This left data from a total of 11,883 probes available for analysis. (supplemental R code #3)

The dbGaP dataset includes expression array data from 849 individuals with a variety of psychiatric diagnoses. After restricting it to schizophrenics and controls it contains 549 individuals and after the elimination of individuals less than 25 years old and individuals whose age is not specified, the cohort consists of 202 schizophrenics and 347 controls (supplemental code #4).

### Identification of differentially expressed transcripts and clustering of schizophrenics

Illumina array probes which detect differentially expressed transcripts were identified using the robust linear mixed effect regression algorithm rlmer{robustlmm} [6] including as fixed effect covariates age, sex, race, and RIN and as a random effect covariate the expression array batch (supplemental code #5). Ingenuity Pathway Analysis (QIAGEN; Redwood City 1700 Seaport Blvd #3, Redwood City, CA 94063) was used to identify pathways containing the differentially expressed genes.

Weighted correlation network analysis (WGCNA) [7] was then used to cluster the schizophrenic patients based on the microarray data for the differentially expressed genes after adjustment for age, sex, race, RIN and expression array batch by linear mixed effect regression analysis (supplemental code #6a). The clustering was validated by perturbation stability analysis (supplemental code #6b) and the demographics of the clusters compared (supplemental code #6c). The R package {igraph} was used to visualize the similarities between the type 1 and type 2 schizophrenics (supplemental code #6d).

Finally, the robust linear mixed effects regression was used a second time, this time to identify the genes differentially expressed in the DLPFC of patients with each of the schizophrenia subtypes analyzed separately (supplemental code #7a and #7b).

### Replication using the CMC RNAseq data

Because of the enormous number of exons represented in the CMC RNAseq dataset and the relatively small number of subjects available (see below), a genome-wide analysis of the RNAseq data was considered to be impractical. Therefore the analysis was restricted to the exons which overlap the Illumina probes which detected differentially expressed transcripts in the NIMH dataset (supplemental code #8a).

The “CMC cohort” is actually three separate distinct cohorts:

1> The University of Pittsburgh ("Pitt") cohort, based on brains specimens from autopsies conducted at the Allegheny County Office of the Medical Examiner.
2> The University of Pennsylvania ("Penn") cohort, based on brain specimens are obtained from the Penn prospective collection and,
3> The Mount Sinai ("MSSM") cohort, based on brain specimens from the Pilgrim Psychiatric Center, collaborating nursing homes, Veteran Affairs Medical Centers and the Suffolk County Medical Examiner’s Office.

As expected, the cohort demographics revealed that the age distribution of the subjects in the Medical Examiner-based “Pitt” cohort is similar to that of the subjects in the Medical Examiner-based NIMH cohort. On the other hand, the subjects in the two primarily hospital-based cohorts, the “Penn” and “MSSM” cohorts, were on the average many decades older (supplemental figure 1; supplemental code #8b). We predicted that the DLPFC transcriptome of young, acutely ill schizophrenics such as those in the “Pitt cohort” would be different from that of older, “burnt out” schizophrenics such as those in the other two cohorts. Preliminary analysis in which the three CMC cohorts were examined separately confirmed that prediction and the analysis reported here is therefore confined to the subjects in the “Pitt cohort”. The exons differentially expressed in the DLPFC of the schizophrenics in that cohort as compared to the controls in that cohort were identified using the R package “edgeR” (supplemental code #8c).

As was done with the NIMH cohort, WGCNA was then used to cluster the schizophrenics in the Pittsburgh-CMC cohort based on the expression levels of the differentially expressed exons. Once again, two subcohorts of schizophrenics were identified with the “type 1” schizophrenics having a DLPFC transcriptome very similar to that of the controls and “type 2” schizophrenics having many differentially expressed transcripts (supplemental code #8d).

## Results

### NIMH Cohort demographics

This cohort is a convenience sample based on Medical Examiner cases for whom the next of kin consented to the use of post-mortem tissue for this purpose. It is, therefore, not necessarily representative of the general population and this study limitation needs to be kept in mind when interpreting these results.

As might be expected in a Medical Examiner cohort where the control subjects include accidental death and homicide victims, men are over-represented in the controls. That imbalance is much more prominent in the Caucasian than African American sub-cohorts (table 1). The cohort is, however, reasonably well balanced in terms of both ethnicity (table 2) and age (figure 1).

**Table 1:** Distribution of schizophrenics and controls by gender

|  | All subjects |  | Caucasians |  | African Americans |  |
| --- | --- | --- | --- | --- | --- | --- |
|  | Controls | Sz | Controls | Sz | Controls | Sz |
| Female | 61 | 72 | 16 | 40 | 43 | 31 |
| Male | 145 | 117 | 76 | 59 | 58 | 54 |

**Table 2:** Distribution of schizophrenics and controls by ethnicity

|  | Control | Sz |
| --- | --- | --- |
| African American | 101 | 85 |
| Caucasian | 92 | 99 |
| Other | 13 | 5 |

**Figure 1:**
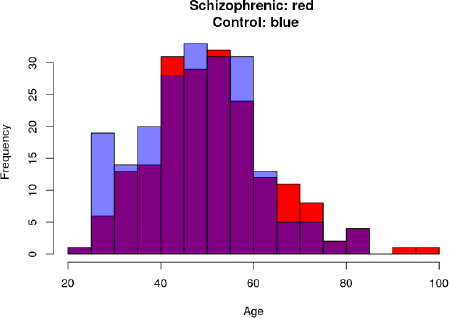
Distribution of schizophrenics and controls by age.

### Genes differentially expressed in schizophrenics as compared to controls

The expression array data included data from 11,883 probes after censoring data from probes which did not detect mRNA in the DLPFC at a statistically significant level or which contained common polymorphisms in the probe sequence.

Robust linear mixed effects regression including RIN, gender, ethnicity, and age as fixed effects and “batch” as a random effect identified 694 array probes which detected transcripts from 633 genes which were differentially expressed in the DLPFC of the schizophrenics at a level of statistical significance which survived Bonferroni-correction. The two genes whose differential expression was most statistically significant were SYNDIG1 (aka TMEM90B, a gene involved in the maturation of excitatory synapses) and PSMB6 (a proteasomal subunit gene), with Bonferroni-corrected P-value less than 10^−15^ for both gene transcripts

Ingenuity pathway analysis identified proteasomal and mitochondrial pathway genes as being overrepresented in the list of differentially expressed genes. The two genes with the largest positive effect size (increased expression in schizophrenics) are MT1X and BAG3, both genes previously identified as being overexpressed in the DLPFC of schizophrenics [8] The gene with the largest negative effect size (decreased expression in schizophrenics) is NPY, a useful marker for specific subclasses of cortical GABAergic interneurons [9–11].

The complete list of the 633 differentially expressed genes is included as on-line supplemental information.

### WGCNA clustering of schizophrenics

In an attempt to identify biologically meaningful subgroups of patients, the expression array data from the schizophrenics for the differentially expressed transcripts was adjusted for the relevant covariates and used as the input to the weighted correlation network algorithm (WGCNA) [7]. That analysis divides the schizophrenics into two groups, “type 1” and “type 2” (figure 2).

**Figure 2:**
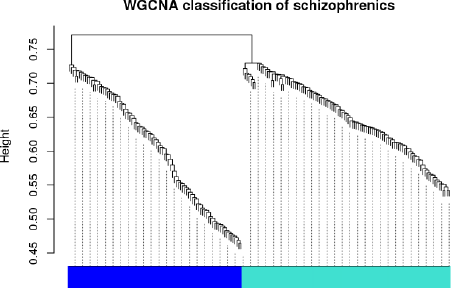
WGCNA clustering of the schizophrenics based on the genes differentially expressed in schizophrenics as compared to controls.

### Perturbation stability of subtype designation

To address the issue of classification robustness, we systematically examined the effect of small changes in the expression array data on subtype classification by repeatedly adding a random error to the covariate-adjusted expression array data used to classify the schizophrenics and tabulating the number of times each schizophrenic was misclassified. The random error introduced into the array data was uniformly distributed over an interval bounded by +/- some fraction of the standard deviation of the data. That fraction is referred to as the “perturbation level” and the subtype classification of any schizophrenic who is misclassified one or more times out of 100 runs was changed from type 1 or type 2 to “intermediate”. So, for example, if after a random error uniformly distributed between -½ and +½ is added to the data a schizophrenic is classified as type 1 once and classified as type 2 the remaining 99 times out of 100, that individual is classified as “intermediate” at a perturbation level of 0.50. Table 3 gives the number of type 1, type 2, and intermediate schizophrenics in this cohort evaluated at several perturbation levels. In everything which follows the schizophrenic subtype designation was made at a perturbation level of 0.50.

**Table 3:** Number of schizophrenics assigned to each subtype at varying levels of random perturbation. (See text for definition of “Perturbation level”)

| **Perturbation level** | **Type 1** | **Type 2** | **Intermediate** |
| --- | --- | --- | --- |
| 0.00 | 92 | 97 | --- |
| 0.05 | 80 | 86 | 23 |
| 0.10 | 80 | 86 | 23 |
| 0.25 | 76 | 85 | 28 |
| 0.50 | 70 | 81 | 38 |

A helpful way to visualize the similarities and differences between the schizophrenics is to examine a family of graphs in which the nodes are individual schizophrenics and edges between schizophrenics are defined as present if the similarity of their DLPFC transcriptomes as measured by their topologic overlap measure (“Rtom” in the WGCNA package) is above some threshold. Taking our lead from the definition of barcodes in topologic data analysis, we systematically varied that threshold and observed how the graph evolved.

**Figure 4:**
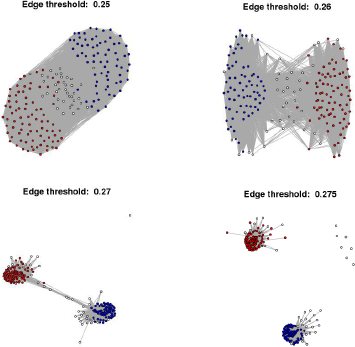
Evolution of the graph of the schizophrenics in the NIMH cohort as the edge threshold is increased from 0.250 to 0.275. In this graph the nodes are individual patients and an edge is defined as present if the topologic overlap measure of their transcriptome similarity (“Rtom” in the WGCNA package) is above the specified edge threshold

As expected, for low values of the threshold the graph has many, many edges and forms a single component. As the threshold is increased the type 1 and type 2 schizophrenics begin to segregate, but the graph remains a single component. At a threshold of around Rtom = 0.275 two connected components form and there are individual isolated nodes. Note however, that at that threshold there are still schizophrenics of ambiguous subtype (classified as “intermediate”) in each of the components. In other words, you could argue that some of the schizophrenics classified as having an intermediate subtype at a robustness threshold of 0.5 should be called, either type 1 or type 2. Excluding them from the type 1 and type 2 subcohorts may be unnecessarily conservative, but preliminary analysis showed that the results described below do not change in any important way no matter how those few individuals are classified.

### Demographics of the type 1 and type 2 schizophrenic cohorts

A comparison of the demographics of the type 1 and type 2 schizophrenics in this study shows that the cohorts are balanced with respect to age, gender, and ethnicity.

**Table 4:** Distribution of type 1 and type 2 schizophrenics by gender

|  | Female | Male |
| --- | --- | --- |
| Type 1 | 31 | 49 |
| Type 2 | 34 | 53 |
| Intermediate type | 7 | 15 |

**Table 5:** Distribution of type 1 and type 2 schizophrenics by ethnicity

|  | African American | Caucasian | Other |
| --- | --- | --- | --- |
| Type 1 | 36 | 42 | 2 |
| Type 2 | 36 | 48 | 3 |
| Intermediate type | 13 | 9 | 0 |

**Figure 5:**
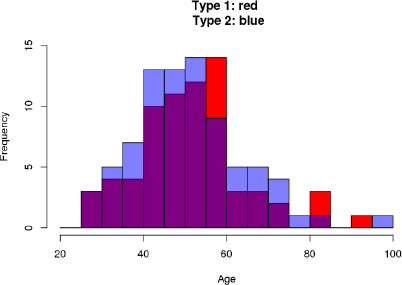
Age distribution of type 1 and type 2 schizophrenics

### Differential gene expression in schizophrenic subtype patients compared to controls

The differential gene expression in the DLPFC is strikingly different in the type 1 and type 2 schizophrenics. There are 3,652 probes to transcripts from 3,200 genes which are differentially expressed in the DLPFC of type 2 schizophrenics at a level of statistical significance which survives Bonferroni correction. On the other hand, there were no differentially expressed transcripts at this level of statistical significance in the DLPFC of the type 1 schizophrenics. This difference in their DLPFC transcriptomes suggests that there is a fundamental biologic difference between these two groups of patients.

Tables 6, 7, and 8 list the genes whose differential expression was most highly statistically significant, the genes with the largest effect size (increased in schizophrenics) and those with the most negative effect size (decreased in type 2 schizophrenics. The complete list of differentially expressed genes is included in the supplemental materials.

### Biologic validation of the distinction between type 1 and type 2 schizophrenia

About half of all schizophrenics, schizoaffective patients, and bipolar patients have what has been described as a “low GABA marker” molecular phenotype based on the expression of GABA neuron markers. Specifically, this subset of schizophrenic patients have reduced expression of GAD67, parvalbumin, somatostatin, and the transcription factor LHX6 in their DLPFC [12, 13].

In this expression array dataset, the Illumina probes for somatostatin and parvalbumin do not detect transcripts at a level significantly different from zero. However, both GAD67 (GAD1) and LHX6 transcripts are detected by the array. In the DLPFC of type 1 schizophrenics there is no statistically significant differential expression of either GAD67 or LHX6 transcripts. In the type 2 schizophrenics however, the P-value <u>after</u> Bonferroni correction for the number of probes on the Illumina array which detect transcripts expressed in the cortex is 5 × 10^−8^ for the differential expression of GAD67; for LHX6 it is 4 × 10^−5^.

In other words, the previously described “low GABA marker” phenotype is highly correlated with type 2 as opposed to type 1 schizophrenia. Since the markers for this phenotype played no role in the distinction between type 1 and type 2 schizophrenics, the differential presence of this phenotype provides a biologic validation of the schizophrenia subtypes.

### The role of RIN

Like many autopsy studies of schizophrenia, the NIMH cohort is slightly unbalanced with respect to RIN, with the DLPFC from schizophrenics having on the average a slightly lower RIN than that from the controls. In this case the mean RIN for the control tissue is 8.1 while that for the schizophrenics is 7.8 (P ≈ 0.01, see figure 6a). Recognizing the potential subtleties involve in properly taking into account variation in RNA quality (see for example [14]) this represents a potential problem for many of these studies. However, in the present study the critical comparison is not between the controls and schizophrenics, but between the type 1 and type 2 schizophrenics. As can be seen in figure 6b, in this study RIN is balanced between those two groups of patients.

**Figure 6a:**
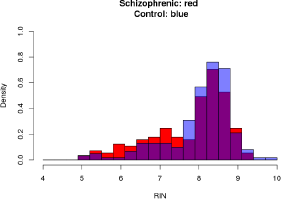
Distribution of RIN in schizophrenics as compared to controls

**Figure 6b:**
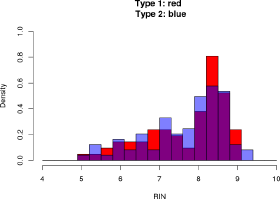
Distribution of RIN in type 1 as compared to type 2 schizophrenics

### The role of neuroleptic therapy

An obvious hypothesis is that the molecular differences between the type 1 and type 2 schizophrenics is due to neuroleptic therapy with the DLPFC transcriptome being normalized in the adequately treated patients (type 1 schizophrenics). Unfortunately there is no information available about the medication compliance of these Medical Examiner patients, but we do have post-mortem toxicology for most of them so we do know which patients had detectable levels of antipsychotics in their blood at death. As can be seen in table 6, there was no statistically significant difference between the type 1 and type 2 schizophrenics in this regard.

**Table 6:** Number of subjects with detectable post-mortem blood levels of neuroleptics. A comparison of the positive vs negative, Type 1 vs Type 2 data is not statistically significant (Chi-squared P ≈ 0.3)

|  | Control | Type 1 Sz | Type 2 Sz |
| --- | --- | --- | --- |
| Negative | 136 | 35 | 28 |
| Not Tested | 70 | 3 | 0 |
| Positive | 0 | 58 | 65 |

### RNAseq replication cohort

As described in the Methods section, the RNAseq data collected by the Common Mind Consortium (CMC) from samples of DLPFC of controls and schizophrenics was used as a replication cohort. In order to have a replication cohort of similar age to the NIMH cohort only the subject from the University of Pittsburgh cohort were included (84 control subjects and 57 schizophrenics). In order minimize the multiple testing problem only data from exons which map to Illumina probes which detected differentially expressed transcripts in the NIMH data were studied. Because many of the Illumina probes map to multiple exons, after censoring exons with less than 10 counts, this resulted in a dataset containing RNAseq data from 3,759 exons.

Of those 3,759 candidate exons, 819 were differentially expressed in the schizophrenic DLPFC at a level of statistical significance which survived Boneferroni correction. WGCNA was then used to cluster the schizophrenics in this cohort based on the RNAseq data from those differentially expressed exons, and once again two subtypes were identified (Figure 7).

**Figure 7:**
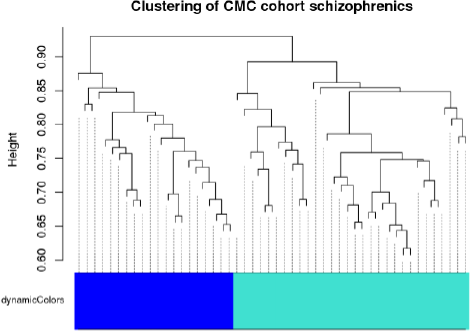
WGCNA clustering of the schizophrenics in the CMC cohort.

The original set of 3,759 candidate exons was then examined for differential expression in the DLPFC of the 23 type 1 schizophrenics or 34 type 2 schizophrenics as compared to the 84 controls. Because of the small number of subjects, rather than use Boneferroni corrected P-values an FDR < 0.05 was used as the criterion for statistical significance. At this level of statistical certainty there were 120 exons differentially expressed in the type 1 schizophrenics, but for the type 2 patients 1,809 of the 3,759 candidate exons were differentially expressed. We interpret these results as replicating those from the study of the NIMH cohort.

## Discussion

This reanalysis of a publicly available expression array dataset identifies 633 genes which are differentially expressed in the dorsolateral prefrontal cortex of schizophrenics as compared to controls at a level of statistical significance which survives Bonferroni correction. More importantly, it demonstrates that schizophrenics can be divided into two molecularly distinct subgroups based on their DLPFC transcriptomes. The “type 1” schizophrenics have a DLPFC transcriptome very similar to that of controls while the “type 2” schizophrenics have a strikingly different DLPFC transcriptome with 3,200 genes differentially expressed as compared to the controls.

A strength of this study is the use of two independent cohorts studied with two different molecular technologies (expression array and RNAseq technology). In addition, there is at least one other published study which, with the benefit of hindsight, reports data consistent with the distinction between type 1 and type 2 schizophrenics. In 2007 Arion et al. used a custom-designed Nimblegen array to measure gene expression levels in the prefrontal cortex of 14 pairs of control and schizophrenic brain samples matched for age, sex and postmortem interval (PMI) [15]. The biclustering of the subjects and genes illustrated in figure 3 of that report shows that 5 of the 14 schizophrenics had frontal lobe transcriptomes similar to the controls, like the type 1 schizophrenics described in this study; the remaining 9 schizophrenics were “type 2” with many differentially expressed genes.

Another strength of the present study is the reliance on robust statistics. Least squares based algorithms are more efficient than the corresponding robust methods IF the data is normally distributed, but they are also exquisitely sensitive to outliers and often give misleading results when the data is from a mixed normal distribution. For a discussion of “regression diagnostics” (the statistical techniques to detect and control for these issues with least squares based algorithms) and robust statistical methods see chapter 6 of Fox and Weisberg and the online appendix “Robust Regression” to that textbook [16] or the documentation for the R package “robustlmm” [6].

This study also takes advantage of graph theory-based analytical methods. Their application here only skims the surface of the opportunities created by the recent advances in applied graph theory and topological data analysis. Because much of that work is being done by mathematicians and computer scientists interested primarily in financial data or computer network security, there is a need for better communication, making the algorithms developed by those analysts available to the biomedical community and vise versa.

Finally, the fact that this study was done on cohorts previously studied by other investigators presents an opportunity to leverage these results and directly apply them to possible re-examinations of those previous studies. For example, the extensive pathway analysis of the CMC RNAseq data by Fromer et al. [2] might be profitably re-examined analyzing the Medical Examiner-based Pittsburgh cohort and the Hospital-based cohorts separately and taking into account schizophrenia subtype. Similarly, the recent study by Tao et al on the expression of alternate GAD1 transcripts in controls and schizophrenics included many subjects in the NIMH cohort [4]. As noted above, GAD1 is one of the genes differentially expressed in the DLPFC of type 2 but not type 1 schizophrenics.

### Scientific implications of these results

#### Increased statistical power and druggable targets for the treatment of schizophrenia

An important aspect of these results is the dramatic increase in statistical power to detect differentially expressed transcripts which is created by analyzing the type 1 and type 2 schizophrenics separately. Only 633 genes were identified as differentially expressed when the data from the type 1 and type 2 patients was combined, but with the recognition of the heterogeneity of that patient population and the analysis of the two subtypes separately that number increased 5-fold to 3,200 genes. An exhaustive review of the molecular biology of those genes and the possible implications of their differential expression in schizophrenic DLPFC is beyond the scope of this report. However, a cursory examination of the list of differentially expressed genes (Supplemental table 2) reveals many druggable targets.

#### A testable hypothesis regarding the pathogenesis of schizophrenia

A common hypothesis regarding the pathogenesis of schizophrenia is that some combination of genetic predisposition and environmental events around the time of birth leads to some persisting alteration in the newborn brain which, although clinically silent itself, predisposes the patient to the development of schizophrenia later in life. From that perspective, the observation that NPY is the most down-regulated gene and that both TAC1 and VIP are high on the list of down-regulated genes in the schizophrenic DLPFC is particularly interesting. Neuropeptide Y (the product of the gene NPY), substance P (produced by proteolytic processing of the TAC1 gene product), and VIP are all well recognized as anatomic markers for particular subsets of inhibitory neocortical interneurons^1^. The expression array and RNAseq data does not address the question of whether there is less of those peptides per interneuron or fewer of those specific types of interneuron, but it does suggest the possibility that the hypothetical perinatal event affects the generation, migration, or maturation of that small subset of the neocortical interneurons.

One might test that hypothesis by counting the number of NPY and TAC1 labeled neurons in the autoradiographic images of schizophrenic and normal DLPFC made public by the Allen Institute. A complimentary approach would be a Nuc-Seq like experiment in which one isolated individual nuclei from DLPFC but replaced the expensive RNAseq with quantitative rtPCR for NPY and TAC1, allowing the study of a large enough sample of nuclei to generate meaningful data regarding these relatively rare interneurons.

#### A testable hypothesis regarding the pathologic neuroanatomy of schizophrenia

The observation that about half of all schizophrenics have a relatively normal DLPFC transcriptome suggests that those (type 1) schizophrenics have physiologically significant pathology elsewhere in their cortex, perhaps in the superior temporal or cingulate gyri. Identifying this hypothetical cortical area where the transcriptome of the type 1, but not type 2 schizophrenics contains many differentially expressed genes would be strong evidence for the physiologic importance of the distinction between type 1 and type 2 schizophrenics and a major step forward in our understanding of the pathobiology of schizophrenia^2^.

Fortunately tissue from both the superior temporal and cingulate gyri from the specific individual patients included in this study is available from the Human Brain Collection Core of the NIMH intramural program. Because the current work provides a list of candidate genes for this next experiment, at least initially the screening of other cortical areas for alterations in the transcriptome of the type 1 schizophrenics could be done as an inexpensive qPCR-based study making this a potentially high-yield, low-cost experiment.

## Acknowledgements

First and foremost we would like to acknowledge the families of the subjects in this study for consenting to the study of this autopsy tissue and for providing clinical information to help establish the psychiatric diagnoses. We would also like to acknowledge the current and former scientific staff of the Intramural Program of the NIMH for collecting the clinical material and expression array data and for making it publicly available on dbGaP. Similarly, we are grateful to the members of the CMC consortium for making their RNAseq data available. CHR is grateful for support from the Henry M. Jackson Foundation and the Center for Neuroscience and Regenerative Medicine at the Uniformed Services University.

## Supplemental Information

R markdown files containing the computer code for the analyses described in this manuscript are available as on-line supplemental files.

A csv file containing a list of the genes differentially expressed in the DLPFC of schizophrenics (without separating the type 1 and type 2 patients) is available as an on-line supplemental file as is a csv file containing list of the genes differentially expressed in the DLPFC of type 2 schizophrenics.

## Footnotes

1 Neuropeptide Y is found in Martinotti cells, neurogliaform neurons, and a subset of the fast-spiking, parvalbumin-positive, basket cells 9. Karagiannis, A., et al., *Classification of NPY-expressing neocortical interneurons*. J Neurosci, 2009. **29**(11): p. 3642-59‥ The first two of those classes of cortical interneurons are well described. The Martinotti cell is a somatostatin-containing interneuron with an axonal plexus in layer 1, making synaptic contact with the spines of pyramidal neuron tuft dendrites. Neurogliaform neuons are non-VIP, 5HTR3A-positive, nitric oxide synthetase-positive neurons with short dendrites spreading radially in all directions and a wider, spherical, very dense axonal plexus. They are present in all layers of the cortex, but are especially prominent in layer 1 where they form the major neuronal component 10. Kubota, Y., *Untangling GABAergic wiring in the cortical microcircuit*. Curr Opin Neurobiol, 2014. **26**: p. 7-14, 11. Tremblay, R., S. Lee, and B. Rudy, *GABAergic Interneurons in the Neocortex: From Cellular Properties to Circuits*. Neuron, 2016. **91**(2): p. 260-92‥ The NPY(+) basket cells are much less well characterized and ignored by many authors. Substance P expression in the neocortex is largely restricted to a specific subclass of basket cells 10. Kubota, Y., *Untangling GABAergic wiring in the cortical microcircuit*. Curr Opin Neurobiol, 2014. **26**: p. 7-14, 11. Tremblay, R., S. Lee, and B. Rudy, *GABAergic Interneurons in the Neocortex: From Cellular Properties to Circuits*. Neuron, 2016. **91**(2): p. 260-92‥ Given the down-regulation of both TAC1 and NPY in the DLPFC of schizophrenics, it is interesting to note that there is a reciprocal interaction between these neurons and the NPY-positive neurogliaform neurons 17. Vruwink, M., et al., *Substance P and nitric oxide signaling in cerebral cortex: anatomical evidence for reciprocal signaling between two classes of interneurons*. J Comp Neurol, 2001. **441**(4): p. 288-301, 18. Kaneko, T., et al., *Morphological and chemical characteristics of substance P receptor-immunoreactive neurons in the rat neocortex*. Neuroscience, 1994. **60**(1): p. 199-211‥ There is, however, an immunohistochemical study using both light- and electron microscopy which describes a second class of large, intensely stained substance P-containing neurons which also express NPY 19. Jones, E.G., et al., *A study of tachykinin-immunoreactive neurons in monkey cerebral cortex*. J Neurosci, 1988. **8**(4): p. 1206-24‥ VIP is found in about 40% of the 5HT3aR expressing interneurons. The majority of these neurons are layer 2/3 bipolar interneurons, but overall they are a heterogenous class of neurons with a variety of morphologies and coexpressed markers 10. Kubota, Y., *Untangling GABAergic wiring in the cortical microcircuit*. Curr Opin Neurobiol, 2014. **26**: p. 7-14, 11. Tremblay, R., S. Lee, and B. Rudy, *GABAergic Interneurons in the Neocortex: From Cellular Properties to Circuits*. Neuron, 2016. **91**(2): p. 260-92‥ Our current understanding of the diversity of cortical interneurons is, however, far from complete and rapid advances in this field are expected with the availability of single-cell and single-nucleus RNAseq technology.

2 If further studies identify a cortical region with transcriptomic abnormalities in the type 1 schizophrenics, it will be important to look for correlations between the clinical features of the schizophrenics and their molecular subtype. For example, if the type 1 patients have molecular pathology in their superior temporal lobes, it would be important to know if those are also the patients with predominantly positive symptoms (including auditory hallucinations).

